# find-tfbs: a tool to identify functional non-coding variants associated with complex human traits using open chromatin maps and phased whole-genome sequences

**DOI:** 10.1101/2020.11.23.394296

**Authors:** Sébastian Méric de Bellefon, Florian Thibord, Paul L. Auer, John Blangero, Zeynep H Coban-Akdemir, James S. Floyd, Myriam Fornage, Jill M. Johnsen, Leslie A. Lange, Joshua P. Lewis, Rasika A. Mathias, Caitlin P. McHugh, Jee-Young Moon, Alex P. Reiner, Adrienne M. Stilp, NHLBI Trans-Omics for Precision Medicine (TOPMed) Consortium, Guillaume Lettre

## Abstract

**Motivation:** Whole-genome DNA sequencing (WGS) enables the discovery of non-coding variants, but tools are lacking to prioritize the subset that functionally impacts human phenotypes. DNA sequence variants that disrupt or create transcription factor binding sites (TFBS) can modulate gene expression. find-tfbs efficiently scans phased WGS in large cohorts to identify and count TFBSs in regulatory sequences. This information can then be used in association testing to find putatively functional non-coding variants associated with complex human diseases or traits.

**Results:** We applied find-tfbs to discover functional non-coding variants associated with hematological traits in the NHLBI Trans-Omics for Precision Medicine (TOPMed) WGS dataset (N_max_=44,709). We identified >2000 associations at *P*<1×10^−9^, implicating specific blood cell-types, transcription factors and causal genes. The vast majority of these associations are captured by variants identified in large genome-wide association studies (GWAS) for blood-cell traits. find-tfbs is computationally efficient and robust, allowing for the rapid identification of non-coding variants associated with multiple human phenotypes in very large sample size.

**Availability:** https://github.com/Helkafen/find-tfbs and https://github.com/Helkafen/find-tfbs-demo

**Supplementary information:** Supplementary data are available.

## 1 Introduction

Genome-wide association studies (GWAS) have identified thousands of common genetic variants (hereby defined as variants with minor allele frequency (MAF) ≥5%) associated with complex human diseases and other quantitative traits. Functional annotation of these variants with maps of open chromatin regions and histone tail modifications has revealed their enrichment within non-coding regulatory sequences (Maurano et al., Vierstra et al.). This suggests that a large fraction of human phenotypic variation is modulated by common variants in sequences that control gene expression. Recently, this hypothesis has been confirmed experimentally in a few robust examples (Musunuru et al., Bauer et al., Lessard et al., Claussnitzer et al.).

In contrast to common variants, most rare genetic variants implicated in human phenotypes have been found in protein-coding exons, mostly because sequencing whole-genomes remained prohibitively expensive until recently. Thus, we still do not know to what extent rare genetic variants in the non-coding human genome can influence inter-individual phenotypic variation. Because rare variants are often missed by the GWAS framework, their identification could yield new loci and genes, or help focus on strong candidate genes at GWAS loci.

Limited statistical power is an important issue for rare variants association testing because of the small number of carriers (by definition) and their very large number in the human genome (Zuk et al., 2014). To lower the multiple hypothesis burden and therefore increase the chance of finding significant associations, analyses of whole-exome sequencing (WES) datasets often collate rare coding variants by genes. For rare non-coding variants found by WGS, collapsing methods based on sliding windows or scanning algorithms have been proposed (e.g. SCANG) (Li et al., 2019; Morrison et al. 2013 and 2017; Natarajan et al., 2018). However, these methods do not consider our current understanding of gene expression regulation, and in particular the important fact that gene expression is controlled by transcription factors (TF) that bind regulatory DNA sequences.

Several algorithms have been developed to predict how DNA sequence variants identified by GWAS can impact the binding of TFs to their corresponding motifs. Tools like atSNP (Zuo et al., 2015; Shin et al., 2019), SNP2TFBS (Kumar et al., 2017) or RSAT (Santana-Garcia et al., 2019) represent powerful computational approaches to calculate TF affinity scores for both alleles at potential regulatory single nucleotide polymorphisms (SNPs). However, these methods were not designed to process WGS data from >100,000s individuals while considering haplotype configurations in order to prioritize regulatory variants for association testing. To address this need, we developed a new tool, find-tfbs, that can efficiently scan a large number of phased WGS and identify genetic variants that create or disrupt transcription factor-binding sites (TFBS) within pre-specified regulatory elements. find-tfbs uses position weight matrices (PWM) to identify and count the number of TFBS occurrences found in each regulatory element of each individual. This information can then be used in standard association testing pipelines. To demonstrate its utility and robustness, we used find-tfbs to analyze associations between TFBS and 15 blood-cell traits in WGS data from 44,709 participants sequenced by the NHLBI TOPMed Project.

## 2 Methods

### 2.1 Scanning phased WGS data to find TFBS

find-tfbs (**Figure 1**) takes three files as inputs: (1) phased WGS data in the BCF format, (2) genomic coordinates of regions of interest, and (3) PWM of prioritized transcription factors. The genomic coordinates file can contain regions identified by open chromatin experiments (e.g. ATAC-seq, DNase1 hypersensitivity), histone tail marks profiling/segmentation or genomic annotation (e.g. gene promoters). It can come from a single cell type or complex tissue. It is also possible to submit multiple coordinates files at once. Since some of these regions overlap, processing time is reduced by merging the overlapping regions, which allows find-tfbs to extract each genetic variant and scan the merged regions only once. At the end of the analyses, find-tfbs dispatches the TFBS it has found and counted in each region to the corresponding cell type/tissue and TF.

**Figure 1.**
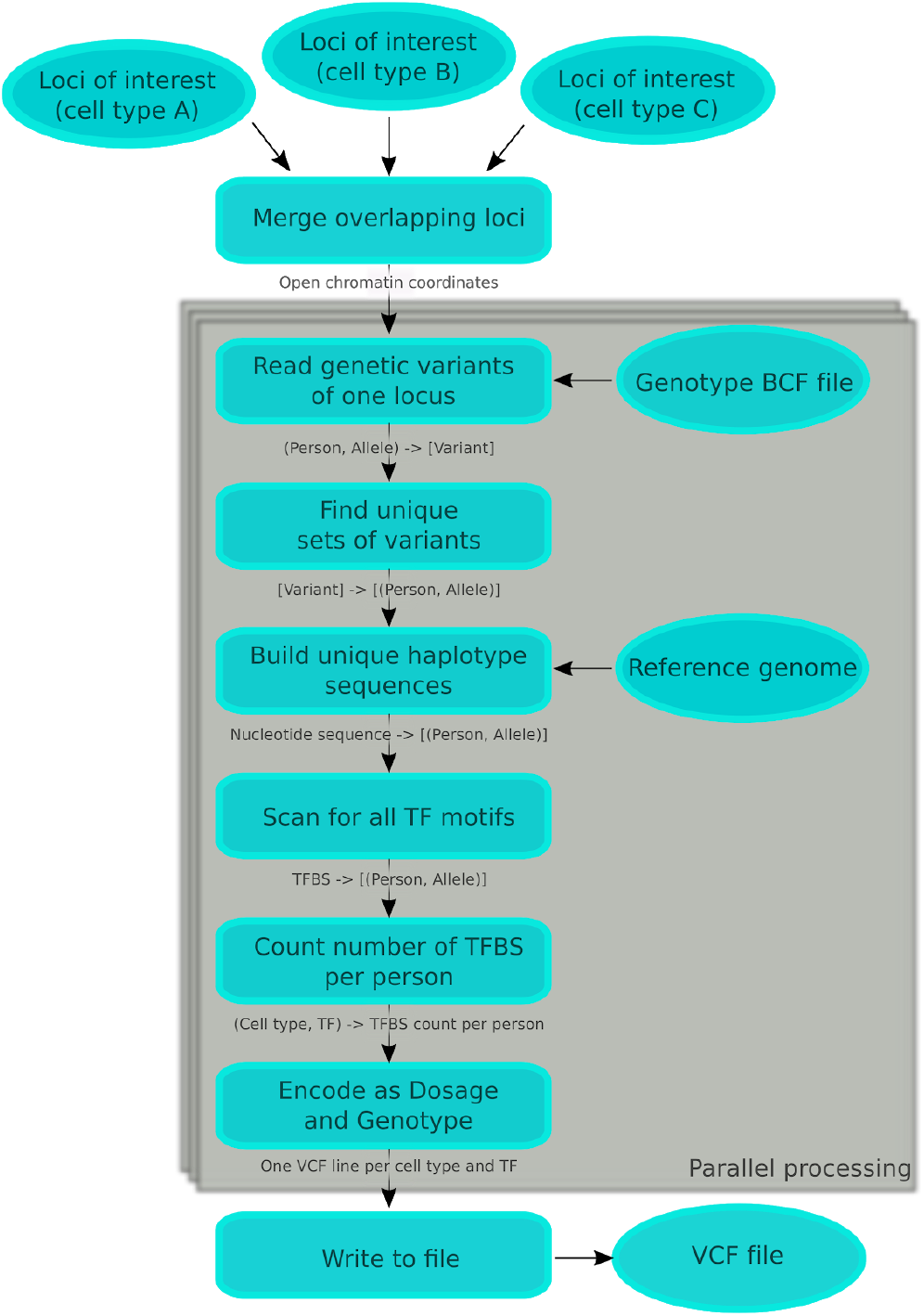
Main processing sequence and data types of find-tfbs. “(X,Y)” is a pair of X and Y: for instance, (Person, Allele) identifies one of the haplotypes of one person. “[X]” is a list of X: for instance, [Variant] contains a list of variants. “X -> Y” represents a hash table with keys of type X and values of type Y, and “TFBS -> [(Person, Allele)]” classifies haplotypes by the binding sites they contain. The keys of a hash table are unique by definition. The sequence within the grey area can be processed independently and in parallel for each locus.

Each merged locus is scanned independently. First, find-tfbs creates a hash table of differences from the reference genome (single nucleotide variants (SNVs) and small insertions-deletions (indels)) in the locus. find-tfbs indexes these differences by haplotype identifiers, where a haplotype identifier represents one strand of one participant. The genetic variants are read from the indexed BCF file. Then, find-tfbs reverses the keys and values of this hash table and uses the reference genome to build the sequence of each unique haplotype: each unique haplotype sequence is associated with a list of haplotype identifiers. This representation is memory efficient: for instance, the haplotype sequence CGGTAACGTGA would exist as a single copy in memory, and it would point to the identifiers (Person5934, Allele2) and (Person4983, Allele1). Using the TF PWMs, find-tfbs scans the unique haplotype sequences for TF motifs on both the forward and reverse strand (Lis et al., 2016; Dai et al., 2007), and the TFBS are associated with a list of haplotype identifiers. Since most variants are rare, the number of distinct haplotypes for a given locus is usually much smaller than 2N (N=cohort size). Scanning unique haplotypes reduces the amount of redundant computing.

Finally, find-tfbs counts the number of TFBS in each cell-/tissue-specific locus for all TFs. find-tfbs uses PWMs and a cutoff scores that are computationally estimated from natural dinucleotide frequencies. In order to minimize the number of false positives, we recommend using the cutoff score corresponding to the most stringent HOCOMOCO standard P-value (0.0001). The result is serialized using the VCF format, and one line is created for each cell type/tissue and transcription factor. find-tfbs discards the lines where the TFBS count frequency (MTCF, defined below) is lower than the threshold (default threshold=0).

### 2.2. Recoding TFBS counts for association testing

First, find-tfbs counts in each participant the number of TFBS found on both alleles for a given region and adds them together. If every individual in the cohort has the same number of TFBS, find-tfbs ignores this region since it is not polymorphic in terms of TFBS count. find-tfbs encodes polymorphic region using the dosage (DS) and genotype (GT) fields of the VCF file format (Danecek et al., 2011). While some association testing pipelines (e.g EPACTS) accept both fields, others only accept the GT field. The DS field accepts any number between 0.0 and 2.0. For our purposes, 0.0 represents the lowest number of TFBS found in a region in the cohort and 2.0 represents the highest number. We interpolate the intermediate TFBS counts and lose no accuracy. The possible values of the GT field are ‘0|0’, ‘0|1’ and ‘1|1’. The lowest and highest numbers of TFBS are encoded as ‘0|0’ and ‘1|1’. The average of the lowest and highest values is encoded as ‘0|1’. Every other value is encoded as the closest encoded neighbour, which makes the GT field less accurate than the DS field for some regions.

### 2.3 Minor TFBS count frequency (MTCF)

For each scanned region and TF, find-tfbs calculates a frequency of TFBS count variations. This allows for simple filtering of regions with too few alternate TFBS counts for association testing. For example, in a cohort of N=100 persons, if 95 individuals have two TFBS in a given region, three participants have one TFBS and two participants have none, then MTCF=(2+3)/N=5%. find-tfbs sums all the groups but the most frequent one. If several groups share the highest frequency, only one of them is taken out of the sum.

### 2.4 Parallelism and performance

Since the merged loci do not overlap (by definition), they can be analyzed separately by any number of worker threads. First, the coordinates of all the loci are sent to a synchronized channel. Each worker thread works in a loop: at the beginning of an iteration, the worker reads the coordinates of one locus, then analyzes it, generates a VCF-formatted result and sends the result to another synchronized channel. The writer thread receives the VCF-formatted strings and writes them sequentially to the result VCF file. Upon completion, the output file handle is flushed and closed by the writer thread. This workflow keeps all the worker threads busy, even if some loci require more processing time than others, and the writer thread guarantees that file writes are sequential. However, the output order is undefined. We save about 200ms of processing time per locus by opening the reference genome file (an indexed FASTA) and the input genotype file (an indexed BCF) at the creation of each worker thread and by keeping the file handles open.

In find-tfbs, the largest data structures are flat arrays. To minimize the number of CPU cache misses and improve performance, they follow the order of the individuals from the input BCF file. These data structures include the ordered list of participants in the input BCF file, the list of participants who share a TFBS or a haplotype sequence, and the number of matches per participant in a locus. None of these data structures are written on disk, in order to increase performance.

### 2.5 Implementation language

find-tfbs is implemented in Rust (Matsakis and Klock, 2014), a programming language that is increasingly used for high performance computing. The explicit memory management of Rust helps the programmer minimize memory allocations and total memory usage, thereby increasing overall performance. Rust provides tools to share data safely in multi-threaded programs, for instance synchronized queues and channels. The language guarantees that any piece of data that is seen by more than one thread can only be accessed safely, which protects the programmer from subtle but common mistakes.

The language also enforces sound error management during compilation. The Rust compiler refuses to compile programs that fail to address several classes of potential runtime errors (e.g memory safety errors, null pointers and uninitialized variables). It has no undefined behavior, unlike C and C++. Rust provides a growing set of bioinformatics libraries. Rust-bio (Köster, 2016) manages FASTA and BED files while rust-htslib (Köster, 2020) manages indexed BCF files.

### 2.6 Application to hematologic traits

A large number of TFs play a role during the proliferation and differentiation of blood cells. However, in many cases, the downstream target genes of these TFs remain unknown. As an example to test find-tfbs, we explored how variation in TFBS counts for 97 TFs modulate 15 blood-cell phenotypes. Phased WGS data and complete blood count (CBC) came from 44,709 participants sequenced by the NHLBI Trans-Omics for Precision Medicine (TOPMed) whole-genome sequencing project, freeze 8 (Taliun et al.). 9870 and 9757 participants have African and Hispanic ancestry, and 25,569 have European ancestry (**Supplementary Table 1**). The TOPMed WGS dataset (freeze 8) is 781 gigabytes after compression.

To prioritize regions more likely to control gene expression, we analyzed open chromatin regions identified in 16 hematopoietic cell types by ATAC-seq (Corces et al., 2016). In our experiments, the list of TFs was based on a literature review, but alternatively it would be possible to perform an exhaustive search with all known TF motifs. When the literature was imprecise, we tested all the relevant progenitor and blood-cell types. For instance, when a source indicated that knockout of a TF was associated with erythrocyte count, we tested this TF in open chromatin regions of erythroblasts and all their available progenitors. When a more precise mechanism was known, we only tested the specific cell type. We restricted the list to the 97 TFs that have a known DNA binding motif in the HOCOMOCO database, version 11 (Kulakovskiy et al., 2018).

We corrected blood-cell traits for age, sex and smoker status by ethnicity and cohort, and then normalized the residuals using inverse normal transformation. We then corrected the normalized phenotypes for population structure within each ethnicity, using the first 10 principal components calculated using 149,454 variants in linkage equilibrium. We used EPACTS for association testing, separately for each ethnicity, using the *q.emmax* algorithm which accounts for cryptic relatedness. Since the variant frequency was already controlled by find-tfbs, we removed the frequency filter in EPACTS, and kept the default values for all other parameters.

Some of the supplementary materials for this experiment can be found in the find-tfbs-demo repository. In particular, the list of relevant transcription factors per cell type and the list of phenotypes per cell type are provided. Genomic coordinates for the open chromatin regions from the different blood cell-types (Corces et al., 2016) are included in the repository for convenience.

### 2.7 Performance

Our experiment was run on a Compute Canada cluster equipped with Intel Xeon Gold 6148 processors. The genotype files occupied a total of 1.006 terabyte for all chromosomes. find-tfbs analyzed 0.41 merged peaks per second on average, using two cores. We used the Linux profiling tool *perf* and observed that loading and decompressing the genotype files was the most resource-intensive task. Building and scanning the unique haplotypes used a relatively small amount of resources.

## 3 Results

### 3.1 Blood-cell trait association results

In this study, we used blood-cell traits to test the implementation of find-tfbs. We arbitrarily defined statistical significance as nominal P-value <1×10^−9^. We acknowledge that this threshold does not rigorously take into account the large number of hypotheses tested and emphasize that association results presented here need to be further replicated. All results that meet this statistical significance threshold are available in **Supplementary Table 2**. The vast majority of the significant associations map to the Duffy/*DARC*, HLA and α-globin loci. Because these loci are already known and genetically complex due to their respective linkage disequilibrium patterns, we did not consider them further in our downstream analyses.

Outside of these three regions, we found 90 combinations of “blood-cell traits/open chromatin regions/TFBS” associated at *P*<1×10^−9^ (**Supplementary Table 2**). This list includes a few highly plausible associations, such as an ATAC-seq peak found in the gene *TAOK1* in megakaryocyte-erythroid progenitor (MEP) cells, which is polymorphic for GATA3 and ETS2 TFBS and associated with mean platelet volume (MPV) in African-ancestry participants. Another interesting association signal, found in European-ancestry individuals, highlights an open chromatin region found in the gene *JMJD1C* in multipotential progenitor (MPP) cells that is associated with platelet counts and include a variable number of binding sites for the TF TWST1, TFE2 and ITF2.

Focusing on associations that map to promoters as annotated in the Ensembl Regulatory Build (Zerbino et al., 2015), we identified signals in the promoters of several genes (**Table 1**). By conditional analyses, we tested if these promoter-based signals were statistically independent from the variants at the same loci that were identified by previous large-scale GWAS for blood-cell traits (Vuckovic et al., Chen et al.). For all but one gene, conditional results were not significant (**Table 1**), suggesting that genetic variants that create or disrupt TFBS in these promoters might explain, at least in part, the GWAS signals. For an ATAC-seq peak in the promoter of *RENBP* located on chromosome X, the association signal remained significant (**Table 1**and **Figure 2**). *RENBP* is an inhibitor in the renin–angiotensin–aldosterone system that regulates arterial blood pressure and it plays an undefined role in early life immune systems (van Bilsen et al.). *RENBP* also serves a catabolic role in sialic acid metabolism (Luchansky et al.). This association signal is present in individuals of African and Hispanic ethnicity and is associated with several red blood cell (RBC) indices (RBC count, RBC distribution width, mean corpuscular volume). This 705-bp open chromatin peak was identified in MEP and encompasses a genetically complex locus: we found 50 distinct haplotypes in the African-ancestry population due to 46 variants (SNPs, indels) and the reference haplotype contains two binding sites for CTCFL, a transcriptional repressor with a similar binding motif to CTCF and that is expressed during spermatogenesis and in certain cancer types (Bergmaier et al.). Out of these 46 variants, two of them disrupt a CTCFL TFBS (X:153946252_G_A, X:153946429_C_T) and one of them creates a CTCFL TFBS (rs7889328)(**Figure 2**). While other variants overlap with the putative binding sites, their individual effects on the PWM scores do not change our model predictions. Further conditional analyses indicated that alleles at rs7889328 accounted for the remaining association signal after controlling for the known GWAS variants at the locus (**Supplementary Table 3**).

**Table 1.**
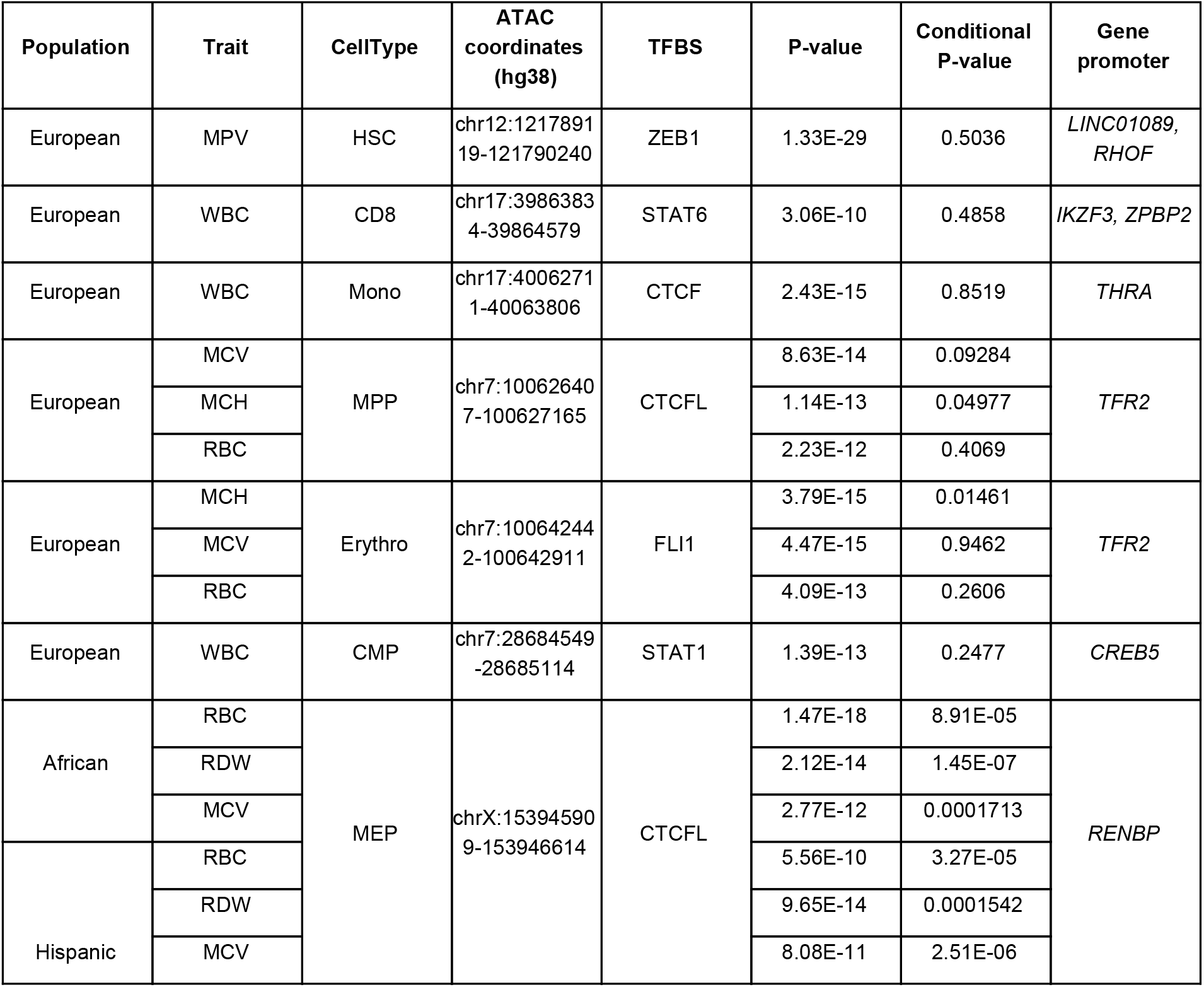
Polymorphic transcription factor binding sites (TFBSs) in gene promoters associate with blood-cell traits. ATAC-seq peaks from different blood-cell types that overlap with ENSEMBL-annotated gene promoters and that include a polymorphic number of TFBS associated with hematological traits. We calculated P-values as described in the **Methods** section; for conditional analyses, we controlled for all genetic variants identified by large blood-cell traits genome-wide association studies located in a 1-Mb window. MPV, mean platelet volume; WBC, white blood cell count; MCV, mean corpuscular volume; MCH, mean corpuscular hemoglobin; RBC, red blood cell count; RDW, RBC distribution width; HSC, hematopoietic stem cell; CD8, CD8+ T lymphocyte; Mono, monocyte; MPP, multipotent progenitor; Erythro, erythroid; CMP, common myeloid progenitor; MEP, megakaryocyte-erythroid progenitor.

**Figure 2.**
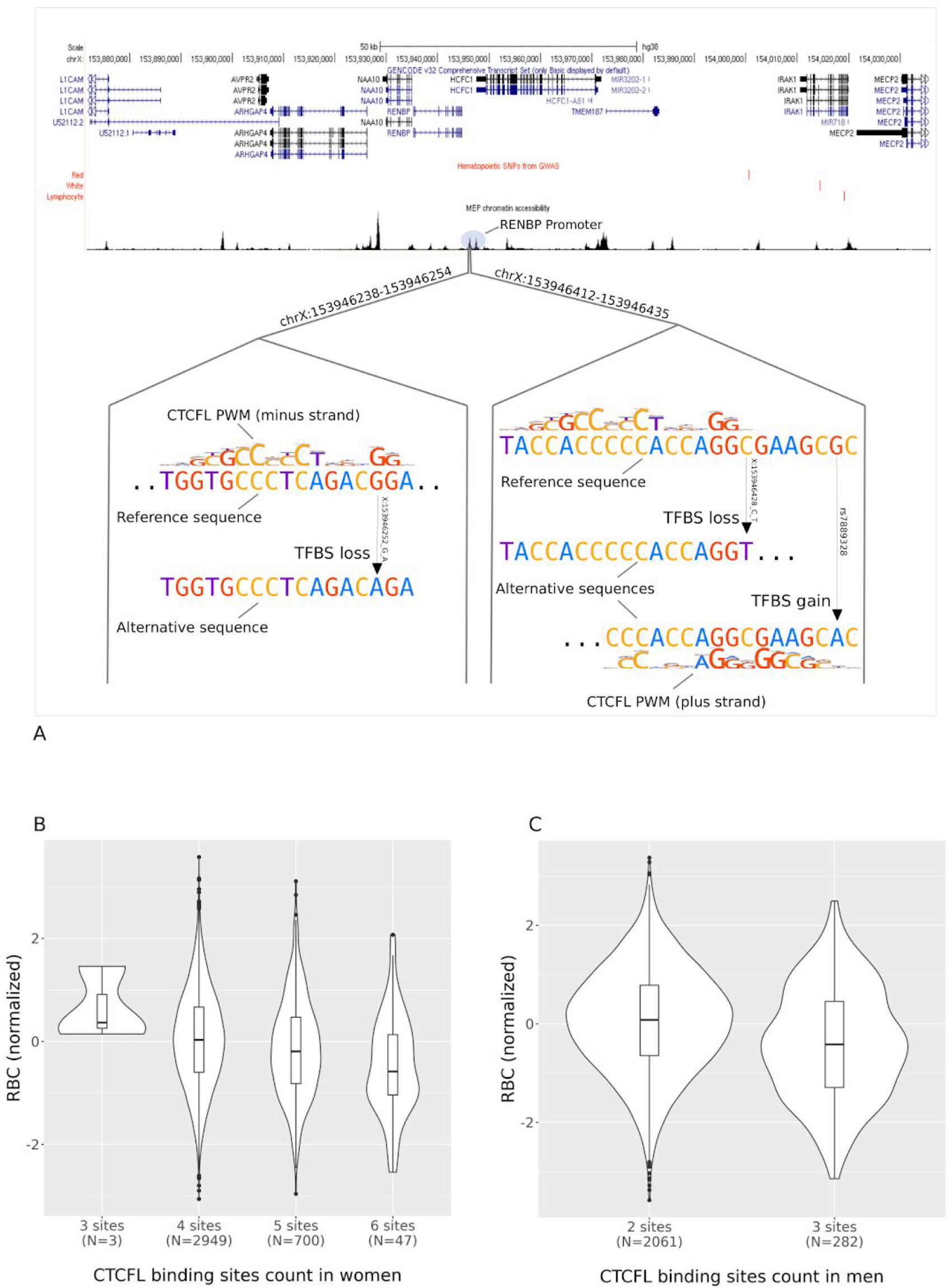
Three genetic variants located in the promoter of *RENBP* and included in an open chromatin region found in megakaryocyte-erythroid progenitor (MEP) cells change the number of CTCFL binding sites and associate with red blood cell (RBC) traits. (**A**) The top panel shows gene annotations at the locus as well as ATAC-seq peaks in MEP. In the bottom panel, we zoom-in two sub-regions in the *RENBP* promoter. The region on the left includes a variant (X:153946252_G_A) where the alternative allele disrupts a CTCFL binding site in the reference sequence (PWM scores: G=−0.188, A=−2.06). The region on the right includes two variants: X:153946428_C_T disrupts a CTCFL motif (PWM scores: C=0.04, T=−0.96) whereas rs7889328 creates a TFBS (PWM scores G=−1.13, A=−0.65). (**B**) Normalized RBC count (y-axis) per number of CTCFL motifs (x-axis) found in the promoter of *RENBP* in women. We summed the number of CTCFL binding sites found in both haplotypes. (**C**) As in (B) but in men. The number of CTCFL motifs is lower in men as RENBP is located on the X-chromosome.

## 4 Conclusion

We developed find-tfbs, a robust algorithm to scan phased WGS data to identify and count TFBS. We tested our tool on the large TOPMed dataset, with an initial focus on hematological traits. We identified many open chromatin regions that harbor genetic variants that create or disrupt TFBS, and that are associated with blood-cell phenotypes. By conditional analyses, we showed that the majority of these associations capture previously identified GWAS loci. Because of our experimental design, such results are interesting because they highlight a possible molecular mechanism. Indeed, find-tfbs combined with association testing tools (e.g. EPACTS) outputs the location of the open chromatin region, the cell-type in which the region was found, and the TF involved, allowing for guided functional characterization of promising GWAS loci.

find-tfbs was purposely designed to be flexible. It will consider all types of genetic variants, including rare variants, and simple plugins can be added to customize the find-tfbs output for other association testing tools. Because of its optimization, find-tfbs can test many phenotypes and regulatory sequence data types in parallel in very large WGS datasets. In the future, considering alternatives to standard PWMs could further improve find-tfbs. For instance, dinucleotide PWMs (Kulakovskiy et al., 2016) and Bayesian Markov models (Siebert and Söding, 2016) outperform mononucleotide PWMs over a variety of datasets by encoding nucleotide correlations. More recently, a promising convolutional neural network called BPNet was able to discover spacing information between motifs, in agreement with known TF-TF interactions (Avsec et al., 2020). As the number and ethnic diversity of WGS data available increase, we expect that find-tfbs will become an extremely useful bioinformatic tool to explore the non-coding regulatory genome implicated in human phenotypic variation.

## Author contributions

S.M.d.B. and G.L. designed the study. All authors contributed data. S.M.d.B. and G.L. wrote the manuscript with contributions from all other authors.

## Acknowledgements

We gratefully acknowledge the studies and participants who provided biological samples and data for TOPMed. The contributions of the investigators of the NHLBI TOPMed Consortium (https://www.nhlbiwgs.org/topmed-banner-authorship) are gratefully acknowledged. The views expressed in this manuscript are those of the authors and do not necessarily represent the views of the National Heart, Lung, and Blood Institute, the National Institutes of Health or the U.S. Department of Health and Human Services.

## Funding

This work has been supported by the Canadian Institutes of Health Research (PJT #168902), the Canada Research Chair Program and the Montreal Heart Institute Foundation (to G.L.). F.T. was supported by the National Heart, Lung, and Blood Institute Division of Intramural Research Funds.

### CHS

Cardiovascular Health Study: This research was supported by contracts HHSN268201200036C, HHSN268200800007C, HHSN268201800001C, N01HC55222, N01HC85079, N01HC85080, N01HC85081, N01HC85082, N01HC85083, N01HC85086, and grants U01HL080295 and U01HL130114 from the National Heart, Lung, and Blood Institute (NHLBI), with additional contribution from the National Institute of Neurological Disorders and Stroke (NINDS). Additional support was provided by R01AG023629 from the National Institute on Aging (NIA). A full list of principal CHS investigators and institutions can be found at CHS-NHLBI.org. The content is solely the responsibility of the authors and does not necessarily represent the official views of the National Institutes of Health.

### SAFS

Collection of the San Antonio Family Study data was supported in part by National Institutes of Health (NIH) grants P01 HL045522, and R01s MH078143, MH078111 and MH083824; and whole genome sequencing of SAFS subjects was supported by U01 DK085524 and R01 HL113323. We are very grateful to the participants of the San Antonio Family Study for their continued involvement in our research programs.

### MESA

WGS for “NHLBI TOPMed: Multi-Ethnic Study of Atherosclerosis (MESA)” (phs001416.v1.p1) was performed at the Broad Institute of MIT and Harvard (3U54HG003067-13S1). MESA is conducted and supported by the National Heart, Lung, and Blood 11 Institute (NHLBI) in collaboration with MESA investigators. Support for MESA is provided by contracts 75N92020D00001, HHSN268201500003I, N01-HC-95159, 75N92020D00005, N01-HC-95160, 75N92020D00002, N01-HC-95161, 75N92020D00003, N01-HC-95162, 75N92020D00006, N01-HC-95163, 75N92020D00004, N01-HC-95164, 75N92020D00007, N01-HC-95165, N01-HC-95166, N01-HC-95167, N01-HC-95168, N01-HC-95169, UL1-TR-000040, UL1-TR-001079, and UL1-TR-001420. Also supported in part by the National Center for Advancing Translational Sciences, CTSI grant UL1TR001881, and the National Institute of Diabetes and Digestive and Kidney Disease Diabetes Research Center (DRC) grant DK063491 to the Southern California Diabetes Endocrinology Research Center.

### HCHS/SOL

Primary funding support to K.N. and colleagues is provided by U01HG007416. Additional support was provided via R01DK101855 and 15GRNT25880008. The Hispanic Community Health Study/Study of Latinos was carried out as a collaborative study supported by contracts from the National Heart, Lung, and Blood Institute (NHLBI) to the University of North Carolina (N01-HC65233), University of Miami (N01-HC65234), Albert Einstein College of Medicine (N01-HC65235), Northwestern University (N01-HC65236), and San Diego State University (N01-HC65237). The following Institutes/Centers/Offices contribute to the HCHS/SOL through a transfer of funds to the NHLBI: National Institute on Minority Health and Health Disparities, National Institute on Deafness and Other Communication Disorders, National Institute of Dental and Craniofacial Research, National Institute of Diabetes and Digestive and Kidney Diseases, National Institute of Neurological Disorders and Stroke, NIH Institution-Office of Dietary Supplements.

### CARDIA

The Coronary Artery Risk Development in Young Adults Study (CARDIA) is supported by contracts HHSN268201800003I, HHSN268201800004I, HHSN268201800005I, HHSN268201800006I, and HHSN268201800007I from the National Heart, Lung, and Blood Institute (NHLBI). CARDIA was also partially supported by the Intramural Research Program of the National Institute on Aging (NIA) and an intra-agency agreement between NIA and NHLBI (AG0005). M.F. is partially supported by U01AG058589 and U01AG052409.

**GeneSTAR** was supported by the National Institutes of Health/National Heart, Lung, and Blood Institute (U01HL72518, HL087698, HL112064) and by a grant from the National Institutes of Health/National Center for Research Resources (M01-RR000052) to the Johns Hopkins General Clinical Research Center.

### GSP

The Genome Sequencing Program (GSP) was funded by the National Human Genome Research Institute (NHGRI), the National Heart, Lung, and Blood Institute 12 (NHLBI), and the National Eye Institute (NEI). The GSP Coordinating Center (U24 HG008956) contributed to cross-program scientific initiatives and provided logistical and general study coordination.

### ARIC

The Atherosclerosis Risk in Communities study has been funded in whole or in part with Federal funds from the National Heart, Lung, and Blood Institute, National Institutes of Health, Department of Health and Human Services (contract numbers HHSN268201700001I, HHSN268201700002I, HHSN268201700003I, HHSN268201700004I and HHSN268201700005I). The authors thank the staff and participants of the ARIC study for their important contributions.

### Amish

The TOPMed component of the Amish Research Program was supported by NIH grants R01 HL121007, U01 HL072515, and R01 AG18728. Email Rhea Cosentino for additional input.

### FHS

The Framingham Heart Study (FHS) acknowledges the support of contracts NO1-HC-25195 and HHSN268201500001I from the National Heart, Lung and Blood Institute and grant supplement R01 HL092577-06S1 for this research. We also acknowledge the dedication of the FHS study participants without whom this research would not be possible.

### JHS

The Jackson Heart Study (JHS) is supported and conducted in collaboration with Jackson State University (HHSN268201300049C and HHSN268201300050C), Tougaloo College (HHSN268201300048C), and the University of Mississippi Medical Center (HHSN268201300046C and HHSN268201300047C) contracts from the National Heart, Lung, and Blood Institute (NHLBI) and the National Institute for Minority Health and Health Disparities (NIMHD). The authors also wish to thank the staffs and participants of the JHS.

### WHI

The WHI program is funded by the National Heart, Lung, and Blood Institute, National Institutes of Health, U.S. Department of Health and Human Services through contracts HHSN268201600018C, HHSN268201600001C, HHSN268201600002C, HHSN268201600003C, and HHSN268201600004C.

### TOPMed Acknowledgements

Whole genome sequencing (WGS) for the Trans-Omics in Precision Medicine (TOPMed) program was supported by the National Heart, Lung and Blood Institute (NHLBI). See the TOPMed Omics Support Table within the Supplementary Document for study-specific omics support information. Core support including centralized genomic read mapping and genotype calling, along with variant quality metrics and filtering were provided by the TOPMed Informatics Research Center (3R01HL-117626-02S1; contract HHSN268201800002I). Core support including phenotype harmonization, data management, sample-identity QC, and general program coordination were provided by the TOPMed Data Coordinating Center (R01HL-120393; U01HL-120393; contract HHSN268201800001I). We gratefully acknowledge the studies and participants who provided biological samples and data for TOPMed.

## Conflict of Interest

The authors declare no conflicts of interest.

## Notes

### Competing Interest Statement

The authors have declared no competing interest.

## References

Avsec,Ž. et al. (2020) Base-resolution models of transcription factor binding reveal soft motif syntax, BioRxiv, doi:10.1101/737981.

Bauer,D. E. et al. (2013) An erythroid enhancer of BCL11A subject to genetic variation determines fetal hemoglobin level. Science (New York, N.Y.), 342(6155), 253–257.

Bergmaier,P., et al. (2018) Choice of binding sites for CTCFL compared to CTCF is driven by chromatin and by sequence preference, Nucleic Acids Res, 46(14), 7097–7107.

Buenrostro,J. (2018) Integrated Single-Cell Analysis Maps the Continuous Regulatory Landscape of Human Hematopoietic Differentiation, Cell. 173, 1535–1548.

Chen,M. H. et al. (2020) Trans-ethnic and Ancestry-Specific Blood-Cell Genetics in 746,667 Individuals from 5 Global Populations. Cell, 182(5), 1198–1213.e14.

Claussnitzer,M. et al. (2015) FTO Obesity Variant Circuitry and Adipocyte Browning in Humans. The New England journal of medicine, 373(10), 895–907.

Corces, M. et al. (2016) Lineage-specific and single-cell chromatin accessibility charts human hematopoiesis and leukemia evolution. Nat Genet 48, 1193–1203.

Dai,X. et al. (2007) A new systematic computational approach to predicting target genes of transcription factors. Nucleic Acids Res, 35, 4433–4440.

Danecek,P. et al. (2011) The variant call format and vcftools. Bioinformatics, 27,2156–2158.

Gate,R et al. (2018) Genetic determinants of co-accessible chromatin regions in activated T cells across humans, Nat. Genet, 50, 1140–1150.

Köster,J. (2016) Rust-Bio: a fast and safe bioinformatics library. Bioinformatics., 32, 444–446.

Kulakovskiy,I. et al. (2018) HOCOMOCO: towards a complete collection of transcription factor binding models for human and mouse via large-scale ChIP-Seq analysis. Nucleic Acids Res, 46, D252--D259.

Kulakovskiy,I. et al. (2016) HOCOMOCO: expansion and enhancement of the collection of transcription factor binding sites models. Nucleic Acids Res, 44, D116--D125.

Kumar,O. et al. (2017) SNP2TFBS - a database of regulatory SNPs affecting predicted transcription factor binding site affinity, Nucleic Acids Res, 45(D1), D139–D144.

Lessard,S. et al. (2017) An erythroid-specific ATP2B4 enhancer mediates red blood cell hydration and malaria susceptibility. The Journal of clinical investigation, 127(8), 3065–3074.

Li,Z et al (2019) Dynamic Scan Procedure for Detecting Rare-Variant Association Regions in Whole-Genome Sequencing Studies, Am. J. Hum. Genet., 104(5), 802–814.

Lis,M and Walther,D. (2016) The orientation of transcription factor binding site motifs in gene promoter regions: does it matter?, BMC Genom, 17, 185.

Luchansky,S. J. et al. (2003) GlcNAc 2-epimerase can serve a catabolic role in sialic acid metabolism. The Journal of biological chemistry, 278(10), 8035–8042.

Matsakis,N. and Klock,F. (2014) The Rust Language. Ada Lett, 34(3), 103–104.

Maurano MT et al. (2012) Systematic localization of common disease-associated variation in regulatory DNA, Science, 337(6099), 1190–1195.

Morrison,AC. et al. (2013) Whole-genome sequence-based analysis of high-density lipoprotein cholesterol. Nat Genet., 45(8), 899–901.

Morrison,AC. et al. (2017) Practical Approaches for Whole-Genome Sequence Analysis of Heart- and Blood-Related Traits. Am J Hum Genet., 100(2), 205–215.

Musunuru,K. et al. (2010) From noncoding variant to phenotype via SORT1 at the 1p13 cholesterol locus. Nature, 466(7307), 714–719.

Natarajan,P. et al. (2018) Deep-coverage whole genome sequences and blood lipids among 16,324 individuals. Nat Commun, 9(1), 3391.

Santana-Garcia,W. et al. (2019) RSAT variation-tools: An accessible and flexible framework to predict the impact of regulatory variants on transcription factor binding, Comput. Struct. Biotechnol. J., 17, 1415–1428.

Siebert,M. and Söding,J. (2016), Bayesian Markov models consistently outperform PWMs at predicting motifs in nucleotide sequences, Nucleic Acids Res, 44, 6055–6069.

Shin,S. et al. (2019) atSNP Search: a web resource for statistically evaluating influence of human genetic variation on transcription factor binding, Bioinformatics, 35(15), 2657–2659.

Sung,M. et al. (2016) Selected heterozygosity at cis-regulatory sequences increases the expression homogeneity of a cell population in humans. Genome Biol, 17, 164.

Taliun,D. et al. (2019) Sequencing of 53,831 diverse genomes from the NHLBI TOPMed Program, BioRxiv, doi:10.1101/563866.

van Bilsen,J. (2020) Seeking Windows of Opportunity to Shape Lifelong Immune Health: A Network-Based Strategy to Predict and Prioritize Markers of Early Life Immune Modulation. Frontiers in immunology, 11, 644.

Vierstra, J. et al. (2020). Global reference mapping of human transcription factor footprints. Nature, 583(7818), 729–736.

Vuckovic,D. et al. (2020) The Polygenic and Monogenic Basis of Blood Traits and Diseases. Cell, 182(5), 1214–1231.e11.

Zerbino,D.R. et al. (2015) The Ensembl Regulatory Build, Genome Biol, 16, 56.

Zuk,O et al. (2014) Searching for missing heritability: Designing rare variant association studies, PNAS, 111(4), E455–E464.

Zuo,C. et al. (2015) atSNP: transcription factor binding affinity testing for regulatory SNP detection, Bioinformatics, 31(20), 3353–3355.

